# Phylogenetic correlation between the Type IV Secretion System and HIP1 suggest an adaptation for horizontal gene transfer conserved at the phylum level

**DOI:** 10.64898/2026.06.24.733823

**Authors:** Ulises Rodriguez-Cruz, Gabriel Moreno-Hagelsieb, Cei Abreu-Goodger, Christian Martínez-Guerrero, Luis Delaye

## Abstract

Most cyanobacterial genomes are rich in the GCGATCGC octamer, also known as Highly Iterated Palindrome 1 (HIP1). Despite its description over three decades ago, the biological function of this highly abundant sequence is only beginning to be elucidated. HIP1 is recognized by two DNA methylases, DmtA and DmtC, and is characterized by its evolutionary conservation and a quasi-periodic distribution within genomes. However, whether the phylogenetic distribution of HIP1 correlates with the presence of functional categories of protein families remains unknown. Here we investigated whether certain protein families share a phylogenetic distribution with this abundant palindromic sequence across cyanobacterial genomes. Our analysis revealed a strong phylogenetic correlation between several proteins of the Type IV secretion system (T4SS) and the abundance of HIP1. This finding aligns with recent discoveries demonstrating that HIP1 enhances DNA transformation in a methylation-dependent manner in two distinct cyanobacterial species. Consequently, we hypothesize that HIP1 function as a conserved adaptation for horizontal gene transfer (HGT) at the phylum level, potentially by serving as a DNA-uptake recognition sequence in cyanobacteria.

**Significance statement:** Scientists have long been baffled by the HIP1 sequence, a short, highly common, repetitive DNA pattern found across almost all cyanobacterial genomes. Our study used a whole-genome evolutionary approach and found that the presence of this repetitive pattern is tightly linked to the presence of a cell’s external DNA uptake system. This tight co-evolutionary relationship suggests that HIP1 isn’t just random genomic feature, but a conserved evolutionary adaptation used by the entire cyanobacterial phylum to specifically enhance their ability to acquire new genes from one another.

## INTRODUCTION

Highly Iterated Palindrome 1 (HIP1), a short octameric sequence (5’-GCGATCGC-3’), was first identified in the cyanobacterium *Synechococcus* PCC 6301 more than three decades ago (Gupta et al. 1993). Despite its early description and remarkable abundance across most cyanobacterial genomes, the functional role of HIP1 has remained largely elusive.

Early research into HIP1 suggested a potential role in genome plasticity and adaptation. This was based on the observation that cadmium tolerance in *Synechococcus* PCC 6301 was driven by the inactivation of the *smtB* locus via the precise excision of a 352-base pair region (Gupta et al. 1993). Because this deletion was flanked by HIP1 sequences, it was proposed that the event occurred through illegitimate recombination between two HIP1 repeats. Subsequent studies demonstrated that HIP1 is the most abundant octameric palindrome in cyanobacteria; however, its polyphyletic distribution suggested that this abundance arose from convergent evolution rather than a single ancestral event (Robinson et al. 1995; Robinson et al. 1997). Furthermore, HIP1 does not introduce insertions or deletions into sequence alignments, leading to the hypothesis that it propagates primarily via nucleotide substitutions rather than insertion events (Robinson et al. 1997).

Nevertheless, identifying a specific molecular function for HIP1 has proven challenging. For instance, Robinson et al. (1997) were unable to detect a HIP1-specific protein complex in *Synechococcus elongatus* PCC 7942 using electrophoretic mobility-shift assays (EMSA). A later study identified an association between HIP1 and a 21-nucleotide sequence named small-dispersed repeat 5 (SDR5) in a few heterocystous cyanobacteria, though the function of SDR5 remains unknown (Elhai et al. 2008). Similarly, direct evidence regarding a role in gene expression has been inconclusive (Vijayan et al. 2011; Xu et al. 2018). While an analysis of 41 cyanobacterial genomes reported a distribution matching the abundance of HIP1 for a version of the glucose 6-phosphate dehydrogenase assembly protein (OpcA), this functional association has not been experimentally tested (Delaye et al. 2011).

Comparative genomic analyses have also revealed that HIP1 motifs are conserved in both coding and non-coding regions, suggesting that their overabundance is maintained by purifying selection (Xu et al. 2018). The observation that HIP1 sites are located at quasi-periodic distances within genomes led to the hypothesis that they play a structural role in the spatial conformation of cyanobacterial chromosomes (Xu et al. 2018).

Recent research indicates that while HIP1 is the most enriched octameric palindrome in cyanobacteria, other highly iterated palindromes (HIPs)–such as GGGATCCC, TCGATCGA, and CAGGCCTG–are also moderately abundant and enriched in certain species (Elhai, 2015; Xu et al. 2018). It has been suggested that cyanobacteria carrying these HIPs possess one or more DNA methyltransferases that recognize these specific palindromic sequences (Elhai, 2015). These findings led to the hypothesis that HIP1 may have originated through a molecular ratchet mechanism involving DNA methylation and a mismatch repair-like (MMR) process mediated by MutS, MutL, and MutH proteins (Elhai, 2015). However, a clear homolog for MutH has not yet been identified in cyanobacterial lineages.

Given that HIP1 contains the recognition sequence for at least two different DNA methyltransferases, a functional association with these enzymes was proposed early on (Robinson et al. 1997; Delaye et al. 2011; Elhai, 2015). In *Synechocystis* sp. PCC 6803, these enzymes are the C5-cytosine methyltransferase DmtC (M.Ssp6803I, *slr0214*) and the N6-adenine-specific DNA methylase DmtA (M.Ssp6803III, *slr1803* or Dam-like). The widespread distribution of DmtC among cyanobacteria rich in HIP1 is particularly notable, especially since the enzyme appears dispensable under standard laboratory growth conditions (Delaye et al. 2011; Elhai, 2015; Hagemann et al. 2018).

The connection between HIP1, methylation, and DNA metabolism was significatively strengthened by findings that DmtC-dependent methylation of HIP1-containing sequences increases the efficiency of exogenous DNA integration into the chromosome of *Synechocystis* sp. PCC 6803 (Wang et al. 2015; Scholz et al. 2019). Further supporting these results, a recent study demonstrated that the inclusion of HIP1 motifs in exogenous DNA significantly increases natural transformation efficiency in a DmtC methylation-dependent manner in *Synechococcus* sp. PCC 7002 (Kamoku and Nielsen 2025).

In summary, despite several independent avenues of research, the primary evolutionary driver of HIP1 accumulation remains to be fully defined. However, the emerging mechanistic link to DNA methylation and transformation strongly points toward a role in competence or DNA uptake. To further explore this possibility, we investigated whether the phylogenetic distribution of HIP1 abundance among cyanobacterial genomes correlates with the presence of specific protein families. We hypothesized that such co-occurrence patterns could reveal the genetic architecture supporting HIP1 utilization, providing new insights into the functional roles of HIP1 and other abundant palindromic octameric sequences in horizontal gene transfer.

## MATERIAL AND METHODS

### Genome selection and phylogenetic inference

All available cyanobacterial genomes classified under the ‘complete_genome’ assembly level were downloaded from the NCBI RefSeq database (https://www.ncbi.nlm.nih.gov/refseq/) on September 5, 2025. This retrieval yielded a dataset of 389 complete and annotated cyanobacterial genomes. The complete list of these genomes, along with their accession numbers, is available in our dedicated GitHub repository (see Data Availability section).

To mitigate taxonomic redundancy, we subsampled the dataset using a phylogenetically informed approach. First, conserved phylogenetic markers were identified using the Genome Taxonomy Database Toolkit (GTDB-Tk) version 1.7.0 (Chaumeil et al. 2020). The list of protein markers is shown in our dedicated GitHub repository. These markers were concatenated and used to infer a Maximum Likelihood (ML) phylogenetic tree with IQ-TREE version 3.0.1 (Minh et al. 2020). The best-fit amino acid substitution model was automatically selected using ModelFinder via the -m MFP option. To partition the dataset into groups of closely related taxa, we applied TreeCluster (Balaban et al. 2019) to the resulting tree of 389-genome phylogeny using a distance threshold (*t*) of 0.05. This clustering yielded 166 distinct groups, 56 of which contained multiple genomes (Table S2). By selecting a single representative genome from each multi-genome cluster, alongside the singletons, we obtained a final non-redundant set of 166 genomes for downstream analyses.

We subsequently inferred the phylogeny of the 166 selected cyanobacterial genomes using IQ-TREE version 3.0.1 (Minh et al. 2020). The best-fit substitution model was automatically selected using ModelFinder (-m MFP). Branch support was assessed with 1,000 replicates of both the Ultrafast Bootstrap (-bb 1000) and the SH-like approximate likelihood ratio test (-alrt 1000). And a second tree was inferred without support values. Topologies from both runs were compared using phylo.io (Robinson et al. 2016), revealing them to be virtually identical, with a normalized Robinson-Foulds (RF) distance of 0, a clade similarity of 100%, and a Euclidean distance of 0.01. Consequently, the phylogeny generated without bootstrap and aLRT support was used for all downstream evolutionary analyses.

The initial phylogeny of all 389 sequences and the final selection are presented in Figure S1 and Figure 2, respectively. All phylogenetic trees were visualized and annotated using the Interactive Tree Of Life (iTOL) v7 (Letunic and Bork, 2024).

### Identification of protein families

Protein families encoded across the 166 non-redundant genomes (including plasmids) were identified using get_homologues.pl version 26012026 (Contreras-Moreira et al. 2013). The software was executed using the OrthoMCL algorithm (-M) with an inflation parameter of 1.5 (-F 1.5), requiring a minimum sequence coverage of 75% (-C 75), and E-value threshold of 1 x 10^-5^ (-E 1e-05), and considering all sequences (-t 0). This analysis generated a presence/absence matrix of protein families across genomes. This matrix was subsequently utilized to identify specific protein families whose genomic distribution correlates with the abundance of the palindromic sequences (see below).

### Abundance of palindromic octamers

We measured the genomic frequency and estimated the observed-to-expected ratio (*O/E*) for a total of all 256 palindromic octamers in the set of all 389 cyanobacterial genomes. To calculate the expected frequency of the octameric palindrome *E*(*W*) within each genome, we employed a second-order Markov model:

Given the octameric palindrome *w_1_w_2_w_3_w_4_w_5_w_6_w_7_w_8_*

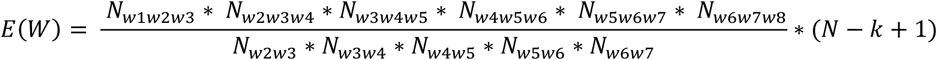

Where: *N_wi…wj_* represents the observed count of the constituent oligonucleotides (trimers in the numerator and dimers in the denominator), *N* denotes the total genome size in base pairs, and *k* is the size of the palindrome (*k* = 8). Scripts used are available in our dedicated GitHub repository.

### Statistical Assessment

To identify palindromic octamers that were significantly overrepresented in each genome, we modeled the occurrence of each sequence using a binomial distribution. The statistical significance of the enrichment was evaluated by calculating the upper-tail probability (*P* [*X* ≥ observed]) under the null hypothesis, by using the pbinom function in R version 4.5.1. For each genome, the number of trials was defined as *N* – *k* + 1 (where *N* is the genome size and *k* = 8), and the probability of success per trial (*p*) was derived from the expected frequency *E*(*W*) estimated by the second-order Markov model (*p* = *E*(*W*) / [*N* – *k* + 1]). To control for Type I error rate arising from multiple testing, the resulting 99,584 *p*-values (encompassing all 256 palindromes across 389 genomes) corrected using the Benjamini-Hochberg False Discovery Rate (FDR) procedure via the p.adjust function in R. Motifs with an adjusted *p*-value < 1 x 10^-308^ were considered significantly enriched.

### Analysis of correlated evolution

To determine whether any protein family exhibited a genomic distribution suggesting correlated evolution with HIP1 overrepresentation, we employed the Discrete method implemented in BayesTraits V5 (Pagel and Meade 2006). This method evaluates whether two binary traits evolve dependently or independently along a phylogenetic tree by comparing the fit of two alternative models: an independent model (where trait transitions are unlinked) and a dependent (correlated) model. The two traits analyzed were: (i) the enrichment status of HIP1 (statistically enriched vs. non-enriched, binarized based on our statistical assessment), and (ii) the presence or absence of each protein family across the 166 cyanobacterial genomes. The analysis was executed using the ScaleTrees option to optimize branch lengths. To ensure convergence, maximum likelihood optimization was performed three times for both the dependent and independent models; the resulting log-likelihood (lnL) values from the three independent runs were averaged for each model. The fit of the two models was formally compared using a Likelihood Ratio Test (LRT) distribution with four degrees of freedom, reflecting the difference in parameter numbers between the dependent (8 parameters) and independent (4 parameters) models. Finally, the resulting P-values were corrected for multiple testing using the Benjamini–Hochberg False Discovery Rate (FDR) procedure as described above.

### Phylogenetic generalized least squares (PGLS) analysis

To complement the discrete trait evaluations, we also modeled the correlation between protein family distribution and the continuous intensity of HIP1 overrepresentation using Phylogenetic Generalized Least Squares (PGLS) regression via the caper package v1.0.4 in R (Orme et al. 2025). For each of the 166 non-redundant cyanobacterial genomes, we quantified HIP1 accumulation through two continuous metrics: (i) the absolute genomic frequency of the octamer (normalized per 1,000 base pairs), and (ii) the observed-to-expected ratio (O/E) derived from our second-order Markov model. To account for differences in scale and ensure equal weight, both variables were standardized by calculating their Z-scores (mean = 0, SD = 1). A composite Selective Intensity Index was then established for each genome as follows:

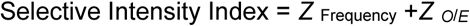

High positive index values identify genomic hotspots characterized by both high absolute abundance and substantial relative enrichment of HIP1. We iteratively fitted PGLS models where the Selective Intensity Index served as the continuous response variable, and the presence or absence of each protein family was treated as a binary predictor. To account for phylogenetic non-independence, Pagel’s lambda (λ) parameter (Pagel, 1999) was simultaneously estimated by Maximum Likelihood (ML) for each model. The resulting *p*-values for the slopes were adjusted for multiple testing using both the Benjamini–Hochberg False Discovery Rate (FDR) and Bonferroni corrections. Protein families with an FDR-adjusted *p*-value < 0.05 were considered to exhibit significant evolutionary correlation with HIP1 intensity.

### Functional Enrichment Analysis

To gain biological insights into the protein families displaying correlated evolutionary patterns with HIP1 intensity and abundance, a functional enrichment analysis was performed using the STRING database v12.0 (Szklarczyk et al. 2023). Representative protein sequences from *Synechococcus elongatus* PCC 7942 corresponding to the significantly correlated families were utilized as queries. Enriched functional categories—from Local Network STRING clusters—were identified using the default network background, with statistical significance set at a False Discovery Rate (FDR) threshold of <0.05. To minimize redundant functional annotations, enriched terms were merged based on a semantic similarity score threshold of ≥ 0.6. Furthermore, to identify discrete functional modules within the resulting interaction network, a Markov Cluster Algorithm (MCL) clustering analysis was implemented with an inflation parameter of 2.

## RESULTS

### Taxonomic representation and dataset composition

All available complete cyanobacterial genomes were retrieved from the NCBI RefSeq database. In total, our dataset encompassed 389 genomes representing 15 of the 20 currently recognized orders of cyanobacteria according to the taxonomic classification proposed by Strunecký et al. (2023). At the family level, the dataset included representatives from 28 of the 55 recognized cyanobacterial families. As shown in Table S1, most of the orders are represented in our dataset and about half of the families. However, the genomic coverage per family is highly skewed; while certain families are densely sampled—such as the Prochlorococcaceae with 143 genomes, the Microcystaceae with 53 or the Thermosynechococcaceae with 33—others are represented by a single genome, like the Anthocerotibacteraceae, the Desertifilaceae or the Gomontiellaceae.

### Frequency and enrichment of palindromic octamers

To evaluate the distribution of palindromic octamers across all these 389 completed cyanobacterial genomes, we analyzed the relationship between genomic frequency and relative enrichment (quantified as the observed-to-expected ratio) for all 256 possible sequences (Figure 1). The Highly Iterated Palindromic 1 (HIP1; GCGATCGC) emerged unequivocally as both the most frequent and the most significantly enriched palindrome across the phylum. Beyond the clear dominance of HIP1, a distinct subset of genomes also exhibited moderately high frequencies and enrichment levels for other specific octameric palindromes. These secondary motifs correspond to previously described variants, such as those identified by Elhai (2015) and Xu et al. (2018).

**Figure 1.**
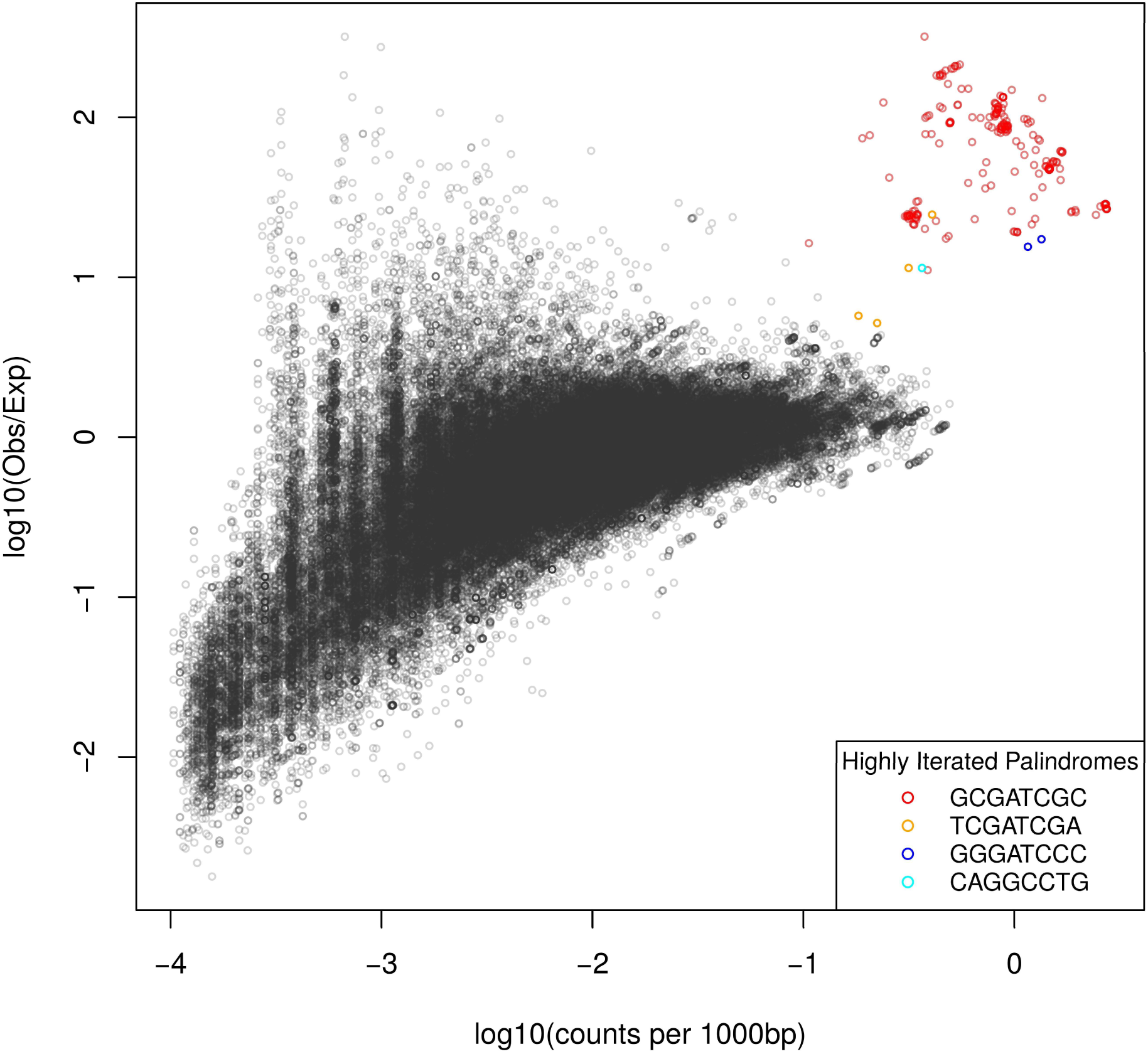
Observed counts and enrichment of octameric palindromes across cyanobacterial genomes. The plot illustrates the relationship between the observed counts and the enrichment of all possible octameric palindromes across all cyanobacterial genomes analysed (N = 389). Enrichment is defined as the ratio of observed to expected frequency. The plot highlights that HIP1 (GCGATCGC) is the most prominent palindrome, exhibiting the highest counts and enrichment. Additionally, the analysis reveals other octameric palindromes that are moderately abundant and enriched in a few specific genomes.

In line with previous reports, several notable evolutionary characteristics emerged regarding these alternative palindromes. First, the octamer TCGATCGA was consistently overrepresented in genomes where HIP1 also exhibited high abundance (Supplementary Material). Second, the CAGGCCTG octamer enriched in *Synechococcus* sp. PCC 6312 was found to harbor a core GGCC motif. This tetranucleotide sequence represents a well-documented palindromic core in cyanobacteria and, much like GATC and CGATGC, functions as a known target for DNA methylation (Elhai 2015; Hagemann et al. 2018). Lastly, the thermophilic hot-spring lineages *Synechococcus* sp. strain JA-2-3B’a(2-13) and *Synechococcus* sp. strain JA-3-3Ab were characterized by a significant enrichment of the alternative HIP variant GGGATCCC, corroborating previous observations by Xu et al. (2018).

### Phylogenetic distribution and exceptions to HIP1 enrichment

To evaluate the evolutionary history of HIP1 enrichment, we analyzed its distribution across a maximum-likelihood phylogenetic tree encompassing all 389 cyanobacterial genomes (Supplementary Figure S1). The phylogeny revealed that HIP1 is significantly enriched across the vast majority of cyanobacterial lineages; however, several notable exceptions emerged. HIP1 overrepresentation is entirely absent in the most diverging orders Gloeobacterales and Thermostichales. Conversely, as noted above, thermophilic hot-spring genomes within the Thermostichales display a significant enrichment of the alternative GGGATCCC variant instead. Following these basal groups, HIP1 became highly abundant in the early-diverging Pseudanabaenales but was subsequently lost across several downstream lineages. Specifically, HIP1 enrichment was conspicuously absent in the vast majority of Synechococcales—particularly within the marine picocyanobacteria clades—with the sole exception of *Synechococcus elongatus* strains. A distinct depletion of HIP1 was also observed in *Synechococcus* sp. PCC 6312 (a member of the Acaryochloridales), where HIP1 is replaced by the CAGGCCTG octamer. Similarly, the motif lacked enrichment in *Leptolyngbya* sp. BL0902 (Nodosilineales) and *Leptolyngbya* sp. NK1-12 (Oculatellales). Finally, HIP1 overrepresentation was notably absent in endosymbiotic cyanobacteria belonging to the Chroococcales where it was lost independently in two different lineages, as well as in their free-living relative, *Synechocystis* sp. LKSZ1. In total, the overabundance of HIP1 was independently lost seven times across the evolutionary history of cyanobacteria.

### Correlated evolution of protein families with HIP1

To investigate whether the presence-or-absence patterns of protein families correlate with the abundance and enrichment of HIP1, we employed the Discrete method implemented in BayesTraits V5 (Pagel and Meade 2006). Briefly, this method assesses the correlated evolution of two binary characters across a phylogenetic tree by evaluating whether transition rates between character states are linked. In our analysis, one binary vector represented the status of HIP1 overrepresentation across genomes, while the other represented the presence or absence of a given protein family.

To mitigate taxonomic redundancy and computational inflation before running the evolutionary models, we subsampled our dataset of genomes using a phylogenetically informative approach. As noted above, the genomic sampling across cyanobacterial families is highly skewed (Table S1). By applying TreeCluster to the initial 389-genome phylogeny with a distance threshold (*t*) of 0.05, we grouped operational taxonomic units (OTUs) exhibiting a phylogenetic distance of 0.05 or less. This clustering yielded 166 distinct groups, 56 of which contained multiple genomes. Selecting a single representative from each multi-genome cluster, alongside the remaining singletons, produced a non-redundant set of 166 genomes (Figure S1 and Table S2). The maximum-likelihood phylogenetic tree of this selected dataset is presented in Figure 2. Crucially, this subset fully preserved the lineages where HIP1 was secondarily lost.

**Figure 2.**
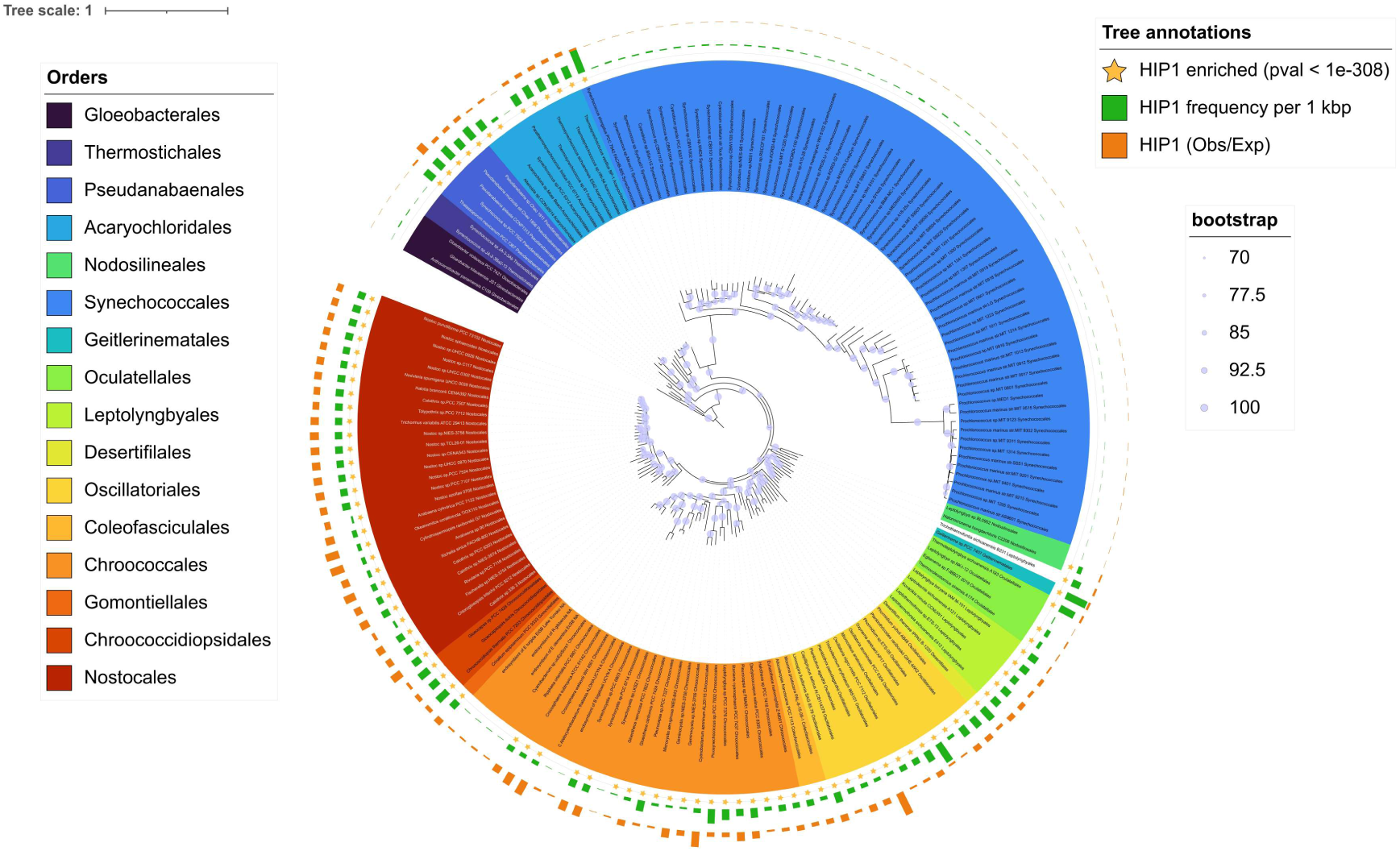
Maximum-likelihood phylogenetic tree of the 166 non-redundant cyanobacterial genomes. External rings indicate the genomic frequency (green bars, normalized per 1,000 bp) and relative enrichment (orange bars, observed-to-expected ratio) of the HIP1 octamer. Yellow stars denote genomes exhibiting a statistically significant overrepresentation of HIP1 (FDR-adjusted *p*-value < 1×10^−308^).

To construct the first binary vector for Discrete analysis, we categorized these 166 selected genomes based on their HIP1 enrichment profiles. Specifically, a genome was assigned a state of ‘1’ if its HIP1 overrepresentation met the stringent statistical threshold of an FDR-adjusted *p*-value < 1×10^−308^; otherwise, it was assigned ‘0’. All colored octamers from Figure 1 met this statistical threshold.

Subsequently, we identified protein families across the 166 non-redundant cyanobacterial genomes using get_homologues.pl, which yielded a total of 105,835 protein families. While, in principle, correlated evolution could have been tested for each of these families against the binary vector of HIP1 overrepresentation, not all pairwise comparisons were expected to be informative a priori. For instance, lineage-specific protein families present in only a single or a few species lack the phylogenetic variance required to detect meaningful correlations with the widespread distribution of the palindrome. Consequently, we restricted the BayesTraits Discrete analysis to protein families present in at least 10 of the 166 genomes. This filtering step narrowed the dataset to a refined set of 8,225 protein families, which were subsequently tested against the binary HIP1 overrepresentation vector.

The Discrete analysis identified 2,148 protein families showing significant correlated evolution with HIP1 overabundance after multiple testing correction (FDR < 0.05). Although statistically significant according to the evolutionary model, many of these protein families did not exhibit a close match with the vector describing HIP1 distribution. In fact, the mean number of mismatches between the HIP1 vector and these 2,148 protein family profiles was 62.8 ± 22.1 (mean ± SD). Therefore, to isolate the protein families displaying the tightest evolutionary co-occurrence with the palindrome, we focused exclusively on candidate families within this statistically significant subset that exhibited the lowest number of profile mismatches.

Mechanistically, two distinct types of mismatches can occur between the reference HIP1 vector and a given protein family vector. A Type A mismatch occurs when a protein family is present in a genome that lacks HIP1 overabundance. Conversely, a Type B mismatch occurs when a protein family is absent from a genome that displays significant HIP1 overabundance. Type A mismatches provide insights into the functional specificity of the relationship between the protein family and HIP1, whereas Type B mismatches reflect its evolutionary necessity. A low number of Type B mismatches suggests that the protein family is nearly ubiquitous wherever HIP1 overabundance occurs. This pattern implies a conserved and potentially essential partnership, as the absence of the protein family appears to be evolutionarily unfavorable in a high-HIP1 genomic background. We reasoned that proteins functionally linked to HIP1 utilization would exhibit fewer Type B than Type A mismatches. This filtering approach effectively prioritizes candidates that are likely integral to HIP1 function, separating them from proteins that are merely broadly conserved across the phylum.

To establish an empirical threshold for permissible Type B mismatches, we evaluated the frequency distribution of these events, which revealed a steady decline from 0 to 5 mismatches (Figure S2). Applying this filter restricted our selection to 110 highly correlated protein families that exhibited at most five Type B mismatches with the HIP1 overrepresentation vector (Table S3; Figure S3).

### Functional enrichment analysis of proteins correlated with HIP1

To evaluate the biological functions overrepresented among the 110 protein families tightly associated with HIP1 overabundance, we performed a functional enrichment and network analysis using the STRING database, a comprehensive resource that compiles records of both physical interactions and functional associations between proteins (Szklarczyk et al. 2023).

By querying representative sequences from *Synechococcus elongatus* PCC 7942 and collapsing redundant categories based on semantic similarity, four clusters exhibited significant functional enrichment (Figure 3, and Tables S3 and S4). Strikingly, the module displaying the strongest enrichment signal was predominantly comprised of core structural components of the Type IV secretion system (T4SS). This was followed by other clusters containing proteins participating in photosynthesis and other processes related to the periplasmic space.

**Figure 3.**
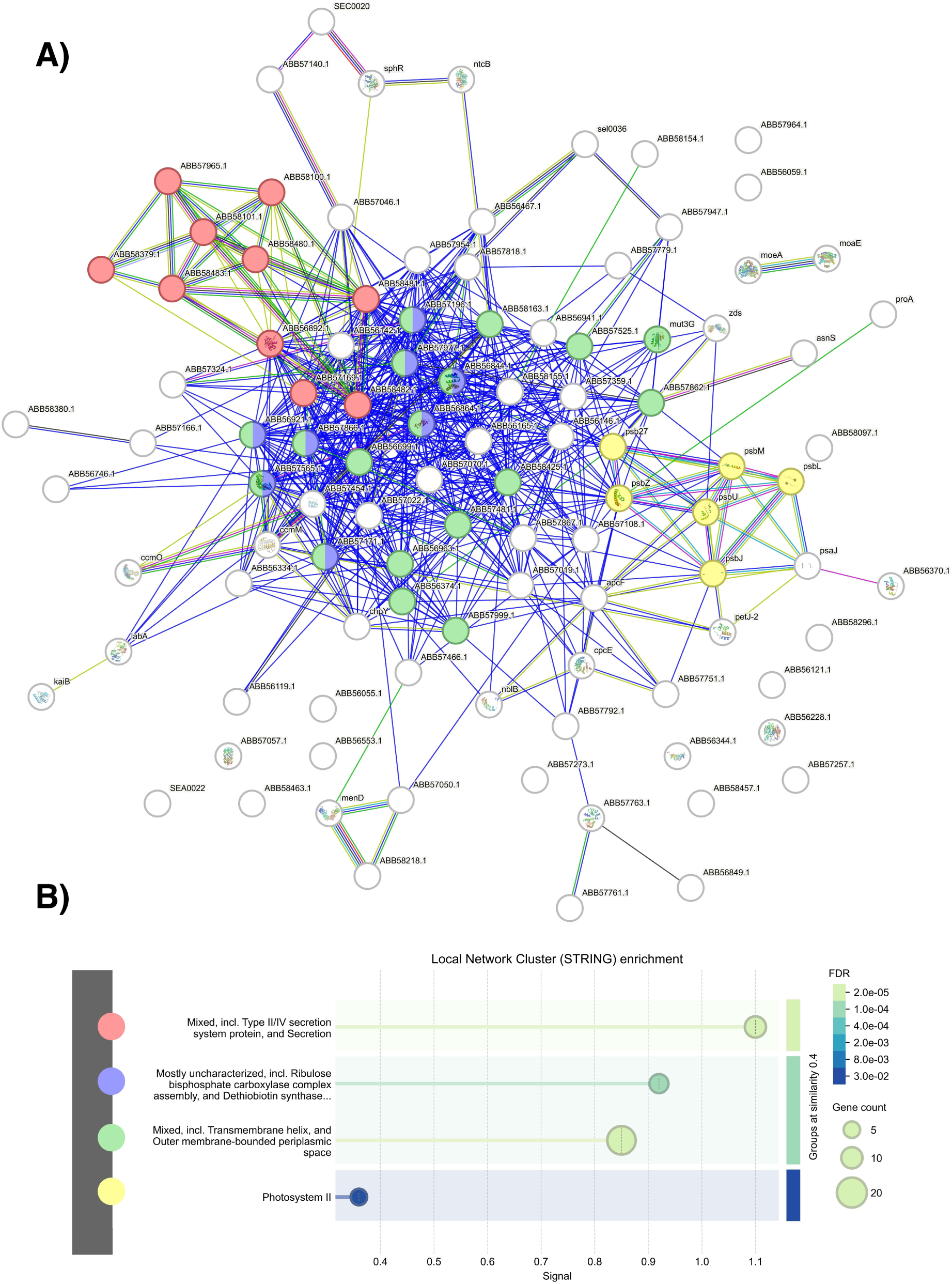
Functional enrichment and interaction network analysis of protein families phylogenetically correlated with HIP1 overabundance. (A) Functional association network generated via the STRING database. Nodes represent protein families, and edges denote independent lines of evidence supporting physical or functional interactions. (B) Statistically significant functional enrichment categories within the network (FDR <0.05). The most prominent enrichment signal corresponds to components of the Type IV secretion system.

### Continuous evolutionary tracking of HIP1 accumulation via PGLS

To transition from a discrete classification of HIP1 overrepresentation to a more nuanced evaluation of its evolutionary dynamics, we mapped the distribution of protein families against the continuous variation in palindrome accumulation across the 166 non-redundant cyanobacterial genomes. While our previous discrete analysis successfully identified broad co-occurrence patterns, it inherently masked differences in the true magnitude of HIP1 enrichment and absolute frequency among genomes. To capture this underlying biological variation, we evaluated a composite Selective Intensity Index—which equally weighs standardized genomic frequencies and observed-to-expected (O/E) ratios—as a continuous response variable against the presence or absence of individual protein families using Phylogenetic Generalized Least Squares (PGLS) regression. By simultaneously estimating Pagel’s lambda (λ) via maximum likelihood, this linear modeling framework effectively disentangled genuine evolutionary correlations from phylogenetic background noise. This high-resolution continuous approach allowed us to identify specific protein families whose genomic presence significantly tracks the precise intensity of HIP1 accumulation, establishing a direct link between genetic architecture and the evolutionary pressure driving palindrome expansion.

Our iterative PGLS linear regressions evaluated the continuous correlation between the composite Selective Intensity Index and the genomic distribution of protein families across the 166 non-redundant cyanobacterial genomes. This analysis identified a total of 99 protein families exhibiting a significant correlation with the intensity of HIP1 overrepresentation at an unadjusted threshold of *p*-value < 0.05. To minimize the inflation of false positives inherent to large-scale genomic screens, we applied stringent multiple-testing corrections. Following this filtering, 14 protein families maintained a robust evolutionary association with an adjusted *p*-value < 0.05 under the conservative Bonferroni correction (Table 1).

**Table 1.** Protein families significantly associated with the continuous intensity of HIP1 overrepresentation identified by PGLS. . For all family clusters, a representative locus tag and protein name from *Synechococcus elongatus* PCC 7942 are shown as references. Statistically significant associations were restricted to families maintaining an adjusted *p*-value <0.05 under the Bonferroni multiple-testing correction. Positive slope values indicate that the presence of the protein family scales with genomic hotspots of high absolute abundance and relative enrichment of the GCGATCGC octamer. Protein families that were also independently captured by the BayesTraits Discrete evolutionary model are marked to highlight cross-methodological convergence.

| LOCUS | STRING protein | Product | p-value | Slope | FDR | Bonferroni |
| --- | --- | --- | --- | --- | --- | --- |
| WP_011242707.1 | ABB57185.1 | Conserved hypothetical protein | 8.30E-11 | 2.23 | 6.62E-07 | 6.62E-07 |
| WP_011244335.1 | ABB58098.1 | Conserved hypothetical protein | 6.98E-09 | 1.57 | 2.78E-05 | 5.57E-05 |
| WP_011244680.1 | ABB57751.1 | PBS lyase; HEAT repeat domain-containing protein | 4.02E-08 | 1.97 | 9.07E-05 | 3.21E-04 |
| WP_011242637.1 | ABB57257.1 | DNA repair protein RadC | 5.31E-08 | 2.17 | 9.07E-05 | 4.24E-04 |
| WP_011378209.1 | nblB | PBS lyase; HEAT repeat domain-containing protein | 5.79E-08 | 2.53 | 9.07E-05 | 4.61E-04 |
| WP_011242445.1 | ccmO | carbon dioxide-concentrating protein CcmO | 6.83E-08 | 2.43 | 9.07E-05 | 5.44E-04 |
| WP_011243311.1 | ABB56553.1 | Pentapeptide repeat-containing protein | 6.45E-07 | 2.24 | 7.35E-04 | 5.14E-03 |
| WP_011242842.1 | ABB57046.1 | Chemotaxis protein CheW | 1.19E-06 | 2.31 | 1.19E-03 | 9.52E-03 |
| WP_011378469.1 | ABB58481.1 | Putative type IV pilus assembly protein PilO | 1.81E-06 | 2.36 | 1.60E-03 | 1.44E-02 |
| WP_011377781.1 | ABB56931.1 | Haloalkane dehalogenase | 2.89E-06 | 1.39 | 2.26E-03 | 2.30E-02 |
| WP_011377872.1 | ABB57155.1 | Putative CheA signal transduction histidine kinases | 3.12E-06 | 1.64 | 2.26E-03 | 2.49E-02 |
| WP_071818121.1 | ABB57964.1 | D-alanyl-D-alanine carboxypeptidase/D-alanyl-D-alanine-endopeptidase | 3.44E-06 | 1.69 | 2.29E-03 | 2.75E-02 |
| WP_231621319.1 | ABB57068.1 | Photosystem II oxygen-evolving complex 23K protein | 4.69E-06 | 2.16 | 2.79E-03 | 3.74E-02 |
| WP_011243128.1 | ABB56746.1 | Cell wall hydrolase/autolysin | 4.91E-06 | 2.20 | 2.79E-03 | 3.91E-02 |

Remarkably, a cross-methodological comparison revealed a high degree of convergence between our independent statistical frameworks: out of the 14 top continuous candidates, nine protein families were also independently identified by the BayesTraits Discrete evolutionary model. This intersection underscores a genuine evolutionary co-occurrence that is independent of the underlying mathematical modeling of the traits.

Among these highly conserved genomic partnerships, several specific functional candidates warrant close inspection. First, the identification of the putative Type IV pilus/secretion system assembly protein PilO (ABB58481.1) provides continuous support for our Discrete network analysis with BayesTraits. This gene family displays a strong positive slope, demonstrating that its presence directly scales with the absolute frequency and relative enrichment hotspots of the HIP1 palindrome. Second, the continuous model highlighted the photo/chemotaxis protein CheW (ABB57046.1) and a putative CheA signal transduction histidine kinase (ABB57155.1). This discovery aligns with our previous Discrete BayesTraits analysis, which independently retrieved a functionally linked CheY-like signal transduction partner, implying that a conserved photo/chemotaxis-like signaling cascade may modulate or co-evolve with HIP1-mediated processes.

Additionally, the continuous screen identified protein families involved in cell envelope maintenance and genetic stability. A positive correlation was observed for a cell wall hydrolase/autolysin (ABB56746.1), suggesting that specific peptidoglycan remodeling processes are evolutionarily favored in high-intensity HIP1 genomic backgrounds, potentially facilitating cell envelope permeability during horizontal gene transfer. Finally, the DNA repair protein RadC (ABB57257.1) emerged as one of the most significant and tightest associations within the dataset. Although the precise molecular mechanism of RadC remains elusive (Attaiech et al. 2008), its close evolutionary tracking of both absolute palindrome abundance and relative overrepresentation points toward a specialized, yet undiscovered, role in HIP1 metabolism.

To further explore the functional connectivity among the top 14 continuous evolutionary candidates, we reconstructed a protein-protein interaction sub-network using the STRING database. Strikingly, this analysis suggested a functional transduction axis connecting five key protein families (Figure S4). Within this network trajectory, the putative CheA signal transduction histidine kinase (ABB57155.1) is directly linked to the photo/chemotaxis protein CheW (ABB57046.1) by several lines of evidence according to STRING database. Crucially, CheW serves as a bridge, likely interacting downstream with a conserved hypothetical protein (ABB58098.1) and the Type IV pilus/secretion system assembly protein PilO (ABB58481.1). Finally, PilO exhibits a direct functional connection with another highly significant conserved hypothetical protein (ABB57185.1). It is important to notice that the evidence supporting these lasts protein interactions is based mostly in gene co-occurrence across genomes. However, the discovery of this coupled interaction chain—spanning from signal transduction components to a core structural protein of the T4SS—suggests that these protein families form a coordinated molecular complex or pathway whose evolutionary maintenance is associated to the intensity of HIP1 genomic accumulation.

## DISCUSSION

### Integration of evolutionary signals: A coordinated genomic network for natural transformation

To obtain a holistic overview of the evolutionary drivers linking protein family distributions with HIP1 dynamics, we synthesized the high-confidence candidates captured by both our discrete (BayesTraits) and continuous (PGLS) phylogenetic frameworks. Taken together, our results point toward a highly coordinated molecular functional network centered on the core structural and components of the T4SS. Specifically, this structural axis encompasses key assembly and pilus factors, including PilO, a pilin-like G/H protein, two PilT paralogs, a PilN-like protein, PilQ, PilM, PilD, and PilB, alongside two conserved hypothetical proteins (ABB56344.1 and ABB56892.1) recently characterized as intrinsic elements of this transport apparatus by Taton et al. (2020). To visualize the spatial and functional layout of these evolutionary associations, we mapped our statistically significant candidates onto a structural model of the *Synechococcus elongatus* PCC 7942 T4SS apparatus (Figure 4).

**Figure 4.**
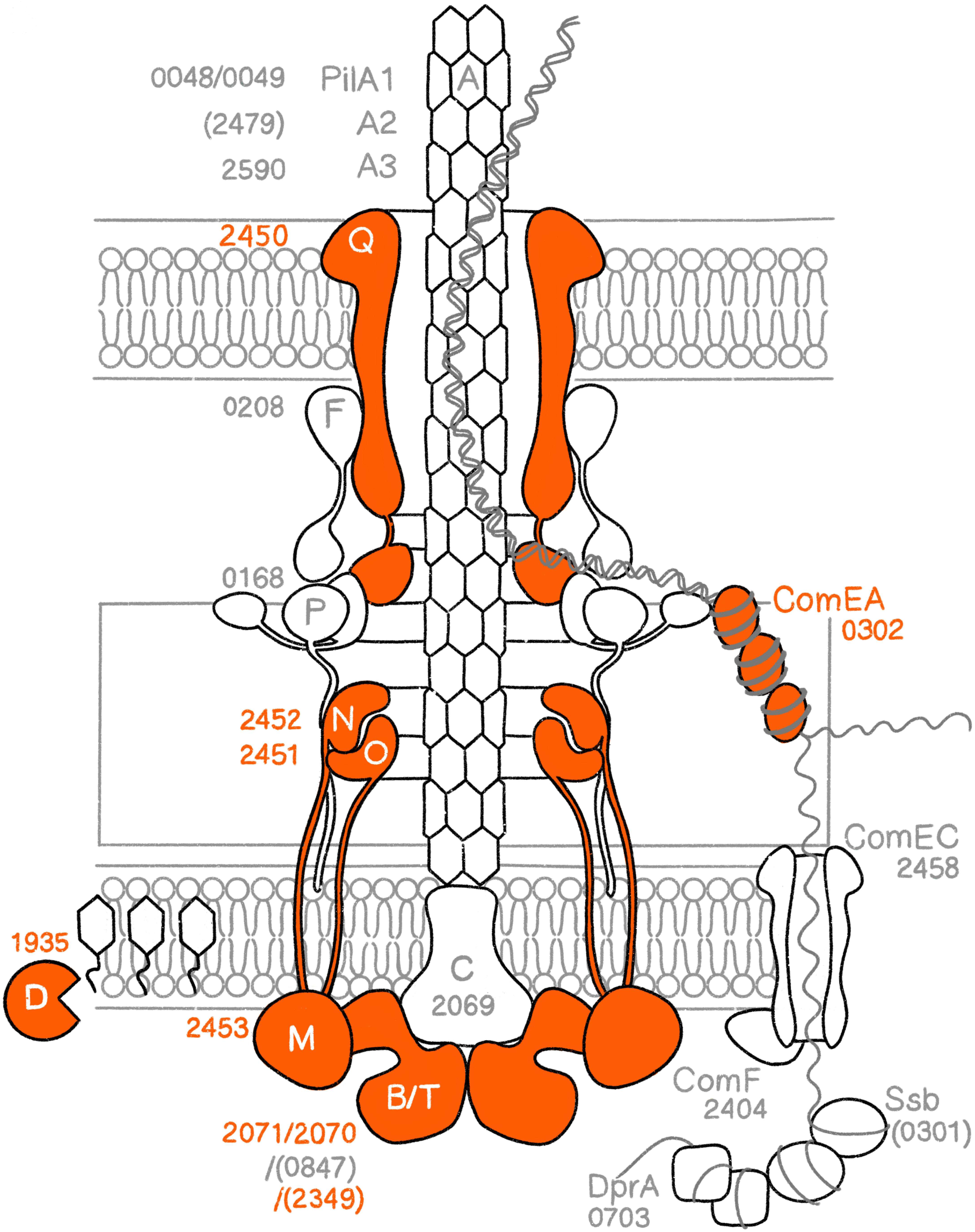
Structural model of the Type IV secretion system (T4SS) in Synechococcus elongatus PCC 7942 highlighting phylogenetically correlated components. The schematic presents the macromolecular architecture of the cyanobacterial T4SS, indicating the cellular localization and predicted functions of its core subunits. Individual proteins are annotated with their respective locus tag numbers. Subunits showing robust evolutionary correlation with HIP1 overabundance across the Discrete and PGLS analyses are colour-marked in orange. Non-essential components for natural transformation under standard laboratory conditions are denoted within parentheses. Modified from Taton et al. (2020).

Beyond this core translocation machinery, our integrated screen successfully identified an array of ancillary protein families that functionally converge on DNA uptake and processing (Table S3). Prominent among these are well-established natural competence factors such as ComEA and ComFB, which operate in close coordination with the T4SS during extracellular DNA binding and internalization (Taton et al. 2020; Samir et al. 2025). Furthermore, the evolutionary signal extended to a photo- and chemotaxis-like signaling cascade composed of CheA, CheW, and an uncharacterized CheY-like protein. Although the molecular function of this specific CheY-like partner remains to be experimentally validated, CheA and CheW—together with an alternative CheY protein not captured in our screen—have been shown to interact directly with the T4SS to control phototactic orientation in Synechocystis sp. PCC 6803 (Han et al. 2024).

Strikingly, this regulatory and structural link is independently supported by functional genomics; co-fitness scores from a Random Barcode Transposon Insertion Site Sequencing (RB-TnSeq) library demonstrate that ComFB operates in the same functional pathway as our CheY-like candidate (Samir et al. 2025). According to our Discrete analysis, this shared fitness cohort also encompasses the hypothetical proteins ABB57466.1 and ABB56892.1, the motility protein HmpF (ABB57169.1)—which is essential for surface pili accumulation (Risser, 2023)—and the core T4SS structural components PilB, PilM, PilN-like, PilO, and PilQ (Table S3). Ultimately, the substantial overlap between the independent datasets of Taton et al. (2020), Samir et al. (2025), and our evolutionary screens provides robust, multi-evidence validation for a unified transformation and motility complex in cyanobacteria.

Our analysis also revealed that this evolutionary co-occurrence spans enzymes involved in cell wall remodeling. These are the D-alanyl-D-alanine carboxypeptidase DacB and the peptidoglycan hydrolase AmiC, which likely facilitate cell envelope permeability during pilus assembly. In addition, the low amplitude/bright protein LabA and the circadian clock protein KaiB which are two key components of the cyanobacterial circadian clock—which has been shown to gate natural transformation in a light-dependent manner (Taton et al. 2020)—were also robustly correlated with HIP1 overrepresentation. Finally, the network is consolidated by essential downstream DNA modification and repair processing factors, including a GIY-YIG family nuclease (ABB57070.1), a recombinase family protein (ABB57196.1), and the elusive DNA repair factor RadC (ABB57257.1). While RadC was initially believed to participate in DNA repair in *Escherichia coli* (Saveson et al. 1999), subsequent studies didn’t support this role in *Streptococcus pneumoniae* (Attaiech et al. 2008). More recently, it was suggested that RadC functions as a nuclease (Iyer et al. 2011). And in cyanobacteria RadC is fused to a conserved N-terminal Helix-hairpin-Helix domain that is often found in proteins involved in DNA replication and repair (Aravind et al. 1999). Additionally, the STRING database shows that RadC (ABB57257.1) interacts with several other proteins involved in DNA repair. Therefore, it is possible that RadC participates in HIP1 metabolism, but further experimental approaches are needed to confirm or discard this hypothesis.

DNA methyltransferases did not reach the rigorous significance threshold required to be included in the final set of protein families showing correlated evolution with HIP1. According to our Discrete analysis, the N6-adenine-specific DNA methylase DmtA exhibited an uncorrected *p*-value of 0.45 and an FDR of 0.51. Conversely, the C5-cytosine methyltransferase DmtC did display a significant uncorrected *p*-value of 0.04; however, it failed to pass multiple-testing correction, yielding an FDR value slightly below 0.10. And the pattern is similar in the continuous PGLS analysis. Interestingly, the markedly higher statistical signal of DmtC relative to DmtA mirrors recent experimental evidence demonstrating that only the former enzyme significantly increases natural transformation efficiencies in cyanobacteria (Wang et al. 2015; Kamoku and Nielsen 2025).

Finally, our analysis also revealed that proteins involved in photosynthesis and other processes show phylogenetic correlation with HIP1. However, despite their statistical significance, the biological relevance of these associations for HIP1 metabolism is not clear. This contrasts with the correlation between proteins participating in T4SS and natural competence, where there is a causal explanation for its phylogenetic association.

### Is HIP1 a universal DNA uptake sequence in cyanobacteria?

Our study reveals a phylogenetic correlation between proteins of the Type IV secretion system and the overabundance of HIP1. The T4SS is known to be involved in the translocation of various substrates, including nucleic acids, across membranes in diverse cyanobacteria (Schuergers et al. 2015; Schirmacher et al. 2020; Chen et al. 2020). This makes their co-evolution with HIP1 particularly intriguing, especially since recent in vitro analyses demonstrated that the inclusion of HIP1 in external DNA significantly increases transformation efficiency in a methylation-dependent manner (Wang et al. 2015; Kamoku et al. 2025).

Environmental DNA offers a cell two primary advantages: it can serve as a source of energy and carbon or be utilized as genetic material for recombination (Mell et al. 2014). A key question is how a cell decides how to utilize the external DNA it encounters. One compelling possibility is that cyanobacteria employ HIP1 as a specific watermark to distinguish DNA originating from other cyanobacteria from that of diverse origins. DNA from another cyanobacteria regardless of the specie is less likely to be harmful and is, therefore, better suited for use as genetic information. Conversely, DNA from a foreign source may be better utilized as a nutrient source.

The hypothesis that HIP1 acts as a universal DNA uptake sequence in cyanobacteria is supported by two lines of evidence. First, our work and that of others show that most cyanobacteria possess all the necessary proteins for a functional T4SS, which is crucial for DNA internalization (Wendt et al. 2019). For instance, recent experimental work showed that natural transformation is not restricted to unicellular species but is also present in *Phormidium lacuna*, a filamentous non-heterocystous cyanobacterium from Oscillatoriales and in *Chlorogloeopsis fritschii* PCC 6912 a member of subsection V cyanobacteria (Nies et al. 2020; Nies et al. 2022).

Second, the observed increase in transformation efficiency mediated by HIP1 in a methylation-dependent manner provides powerful functional evidence. This effect has been shown in two phylogenetically distinct species: *Synechococcus* sp. PCC 7002 (order Chroococcales, subclass Oscillatoriophycideae) and *Synechocystis* sp. PCC 6803 (order Synechococcales, class Cyanophyceae). These species are separated by millions of years of evolution (Demoulin et al. 2019). The functional conservation of HIP1 in such distantly related organisms supports the hypothesis that it acts as a broadly utilized DNA uptake sequence. If this is the case, HIP1 would represent a significant adaptation for horizontal gene transfer (HGT) specifically adapted for inter-species DNA exchange within the cyanobacterial phylum. The higher proportion of phylogenetic conflicts within cyanobacteria than between cyanobacteria and other phyla supports this hypothesis (Zhaxybayeva et al. 2006).

## CONCLUSIONS

This study reveals a phylogenetic correlation between T4SS proteins and the highly iterated palindrome HIP1. This co-evolutionary relationship is supported by recent studies demonstrating that HIP1 enhances DNA transformation efficiency in a methylation-dependent manner. Given that this effect has been demonstrated in two distinct cyanobacterial species belonging to different orders, and that both the T4SS and HIP1 are widely conserved across the phylum, we propose that HIP1 serves as a conserved adaptation for HGT within cyanobacteria.

## DATA AVAILABILITY

Phylogenetic trees in *newick* format, frequency and abundance of all palindromic octamers in cyanobacterial genomes, the vectors used for the *Discrete* analysis in BayesTraitsV5, the version of the genomes used in this analysis and the code in Perl and R can be found at https://github.com/luisdelaye/HIP1_associated_proteins.

## SUPPLEMENTARY DATA

Supplementary Data are available online.

## ACKNOWLEDGEMENT

During the preparation of this manuscript, the author(s) used Artificial Intelligence solely for improving the grammar and refining the clarity and readability of the manuscript. The authors confirm that they have reviewed, edited, and verified the content as needed and take full responsibility for the content of the publication.

## Contributions

L.D. conceived and led the study, performed the bioinformatic analysis, wrote the first draft of the paper, and supervised U.R.C. and C.M.G. U.R.C. performed bioinformatic analyses. G.M. discussed the study with L.D. and provided suggestions for the bioinformatic analysis. C.A.G. discussed the study with L.D. and advised on statistical and bioinformatic analysis. C.M.G. performed bioinformatic analyses and helped with the cluster facilities from Laicbio. All authors read and approved the final manuscript.

