## Supplementary Figures S1, S2, S3 and S4 for "Phylogenetic correlation between the Type IV Secretion System and HIP1 suggest an adaptation for horizontal gene transfer conserved at the phylum level"

**Supplementary material.**

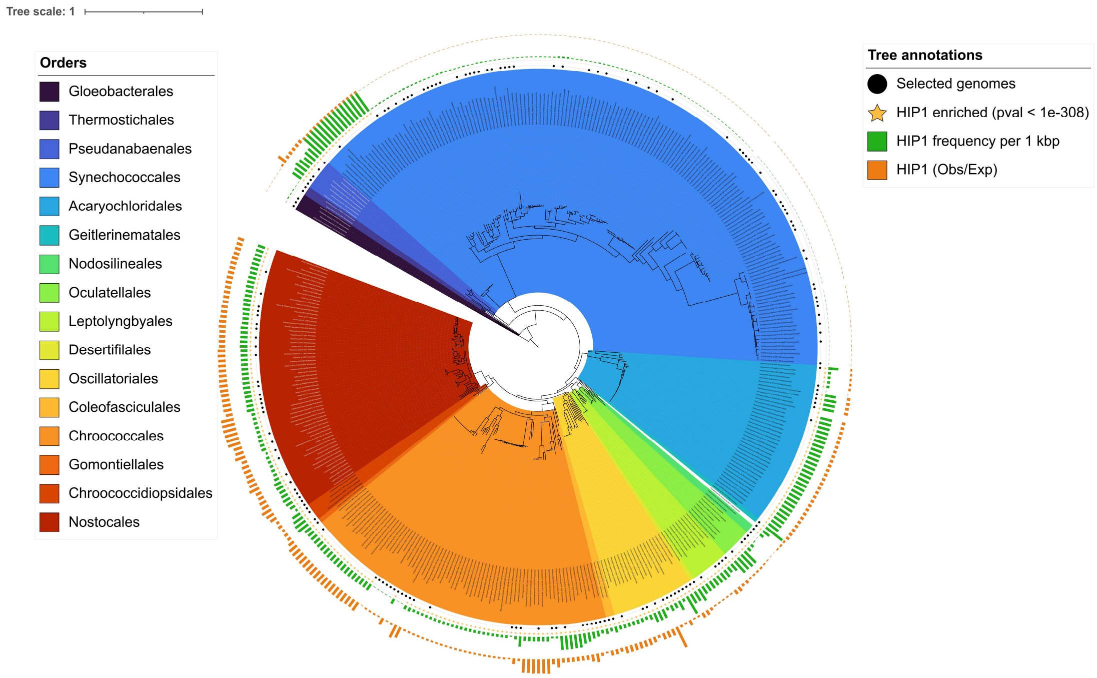


**Figure S1. Maximum-likelihood phylogenetic tree of 389 cyanobacterial genomes**. External rings indicate the enrichment (orange bars) and genomic frequency (green bars) of the HIP1 octamer. Yellow stars denote genomes with a statistically significant enrichment of HIP1. Black dots indicate the 166 representative genomes selected for downstream analyses using the TreeCluster dereplication approach.

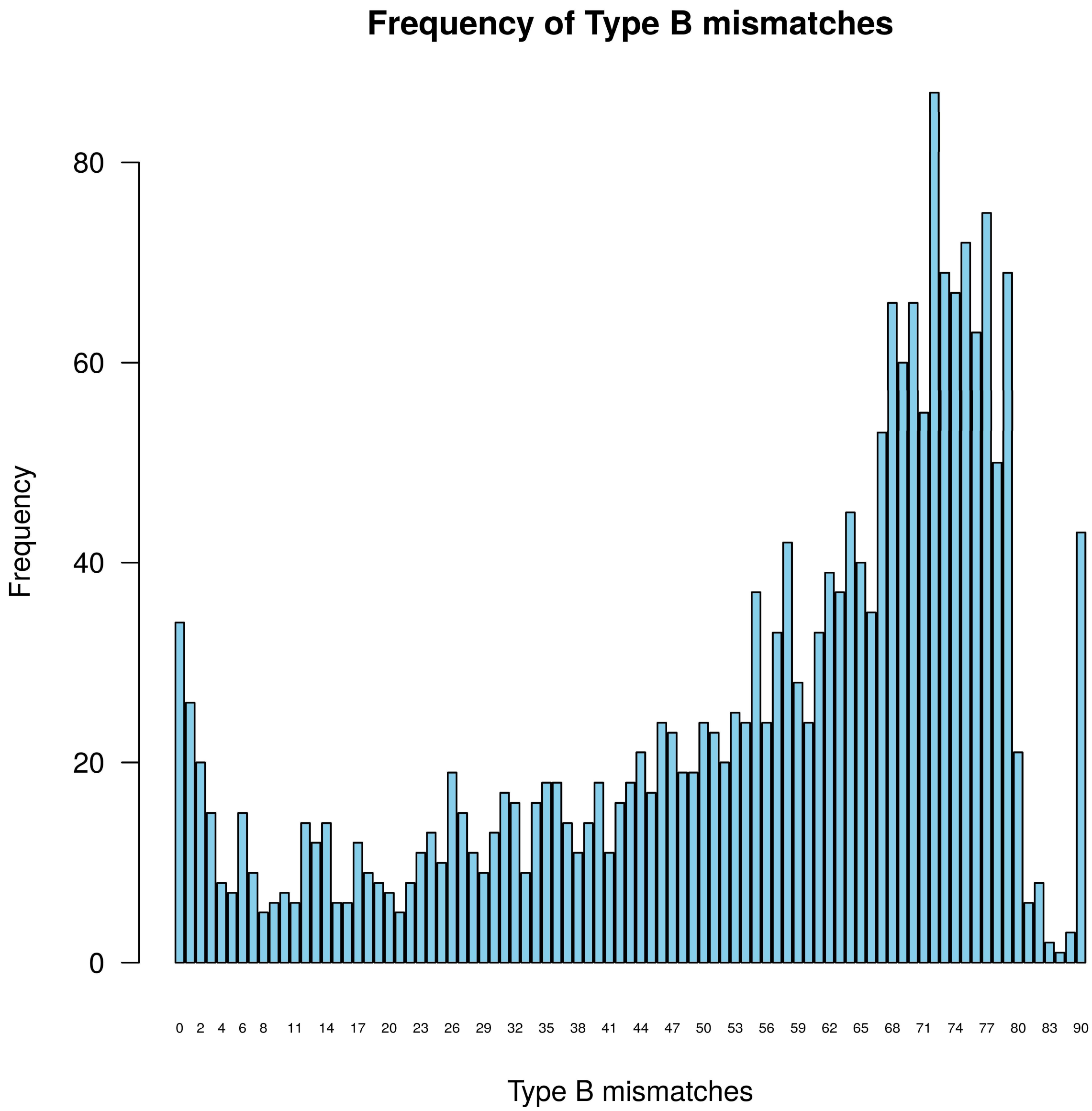


**Figure S2. Distribution of Type B mismatches among statistically significant protein families**. The histogram displays the genomic frequency of Type B mismatches for the subset of 2,148 protein families that exhibited significant correlated evolution with HIP1 overabundance (FDR < 0.05) in the BayesTraits Discrete analysis. The empirical threshold was established at a maximum of five Type B mismatches, capturing the initial high-frequency distribution following a steady decline before the frequency increases again at six mismatches.

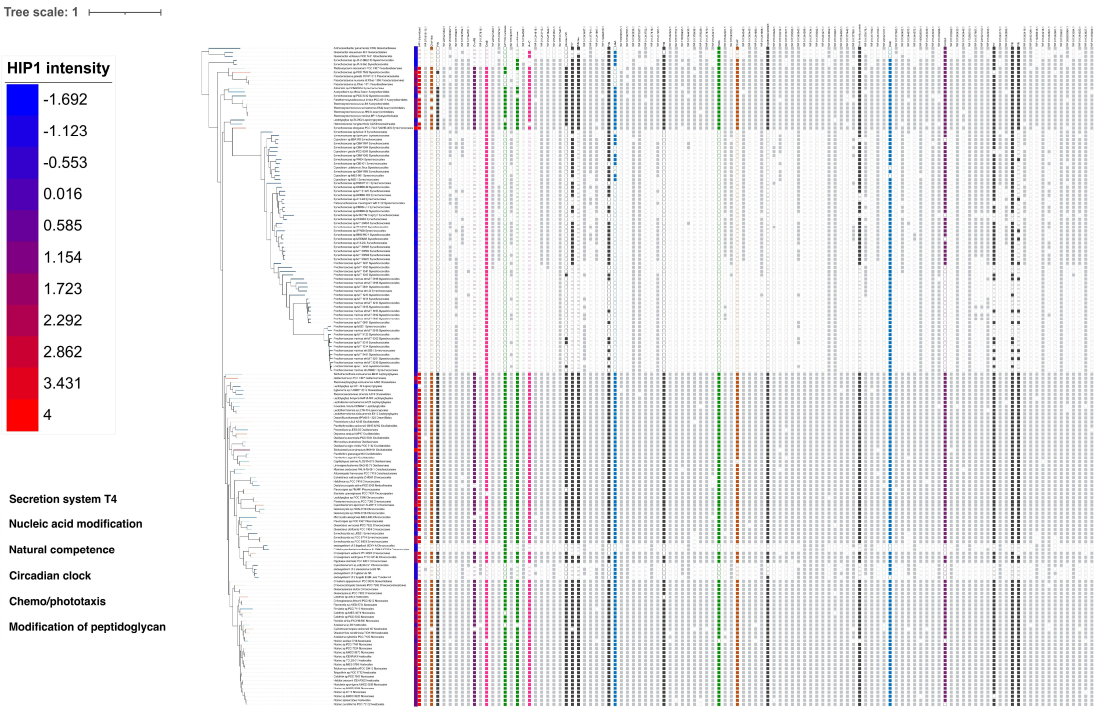


**Figure S3. Phylogenetic distribution of highly correlated protein families across the non-redundant cyanobacterial dataset.** Maximum-likelihood phylogenetic tree of the 166 non-redundant cyanobacterial genomes mapped against the presence/absence profiles of the 110 protein families selected after multi-step filtering. Candidate families were restricted to those displaying robust correlated evolution with HIP1 overabundance in the BayesTraits Discrete analysis (FDR<0.05) and exhibiting an empirical threshold of at most five Type B mismatches relative to the reference HIP1 vector. Matrix columns represent individual protein families, where coloured cells indicate presence and white cells denote absence. Specific functionally relevant protein families discussed in the main text are highlighted with distinct colour coding [e.g., black for type IV secretion system components] to facilitate evolutionary tracking across the phylum.

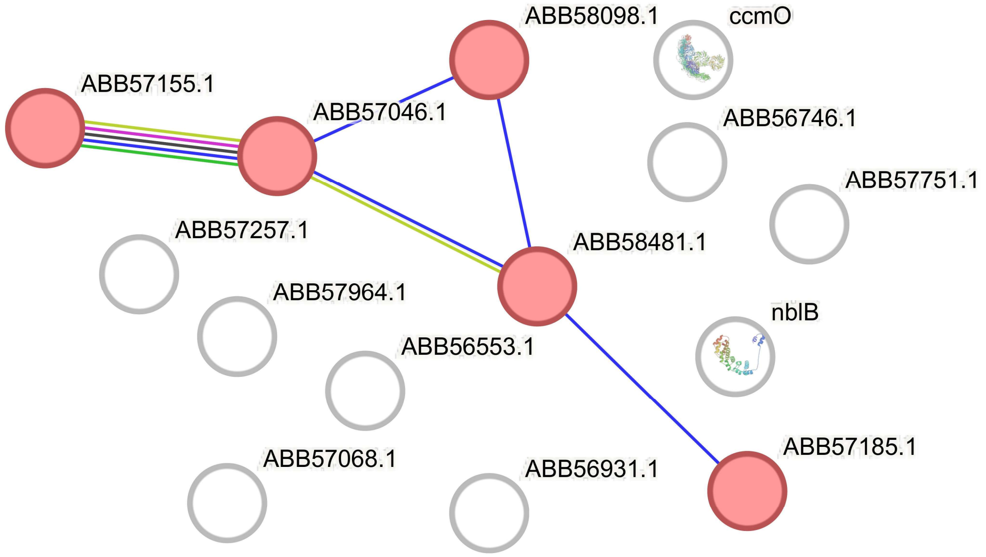


**Figure S4. Characterization of a conserved protein interaction axis among top PGLS candidates.** The network diagram highlights a continuous functional and physical interaction pathway retrieved via the STRING database among the high-confidence protein families identified by the continuous PGLS regression. Nodes represent specific protein families from *Synechococcus elongatus* PCC 7942, and edges denote independent lines of experimental or genomic evidence supporting functional co-occurrence. The topological layout illustrates a direct transduction chain running from chemotaxis and signal transduction elements (CheA/ABB57155.1 and CheW/ABB57046.1) through conserved hypothetical protein (ABB58098.1) and the core Type IV secretion system assembly protein PilO (ABB58481.1), culminating in another hypothetical protein (ABB57185.1).
